# Structural and Functional Mechanisms Underlying Activation Gate Dynamics and IFM Motif Accessibility in Human Na_v_1.5

**DOI:** 10.1101/2025.06.29.662239

**Authors:** Rupam Biswas, Ana Laura López-Serrano, Apoorva Purohit, Angelina Ramirez-Navarro, Hsiang-Ling Huang, Xiaolin Cheng, Sarah M. Heissler, Isabelle Deschênes, Krishna Chinthalapudi

## Abstract

Voltage-gated sodium channels are vital for regulating excitability in muscle and nerve cells, and their dysregulation is linked to a range of diseases. However, therapeutic targeting of Na_v_ channels remains challenging due to a limited understanding of their gating mechanisms. Here, we present a cryo-EM structure of human Na_v_1.5 in an intermediate open state, stabilized by interactions between the N-terminal domain and the S6_I_ segment. This structure reveals a previously uncharacterized Na^+^ binding site adjacent to the conserved inactivation (IFM) motif. Molecular dynamics simulations demonstrate that monovalent cations stably occupy this site, while electrophysiological recordings demonstrate that ion binding modulates IFM motif docking and fast inactivation kinetics. Our findings reveal that IFM accessibility is dynamically regulated in this intermediate state, challenging the canonical hinged-lid model of fast inactivation. Collectively, our study provides a revised structural framework for Na_v_1.5 gating mechanisms, suggesting an alternative pathway for ion accessibility that may inform better mechanistic and therapeutic strategies for treating Na_v_1.5-related cardiac arrhythmias.

## Introduction

Voltage-gated sodium channels are fundamental to the initiation and propagation of action potentials. In the heart, the predominant sodium channel isoform Na_v_1.5 (*SCN5A*) is indispensable for the rapid depolarization phase of the cardiac action potential^1^. Upon membrane depolarization, Na_v_1.5 channels activate with sub-millisecond kinetics, permitting a transient inward sodium current (I_Na_) that drives the rapid upstroke of the action potential^2^. Channel opening is rapidly followed by fast inactivation, a tightly coupled process that closes the pore and terminates sodium influx within milliseconds^3^. Proper coordination of activation and inactivation is essential for maintaining normal cardiac electrical activity^4^. Perturbations in Na_v_1.5 function, such as mutations that impair fast inactivation, are implicated in a spectrum of inherited and acquired cardiac arrhythmias^5-7^. Thus, the precise regulation of Na_v_1.5 gating transitions is essential for maintaining normal cardiac excitability and rhythm.

Na_v_1.5 is composed of a large pore-forming α-subunit organized into four homologous domains (D_I_ to D_IV_), each comprising six transmembrane segments (S1-S6) arranged in a pseudo-tetrameric configuration^8^. Within each domain, the S1-S4 helices form the voltage-sensing domain (VSD), whereas the S5-S6 helices and the intervening reentrant loop constitute the pore domain (PD)^8-10^. The activation gate (AG) is formed by a cluster of hydrophobic residues that line the intracellular ends of the S6 helices^9,10^. Fast inactivation is governed by the highly conserved isoleucine-phenylalanine-methionine (IFM) motif, also known as the inactivation particle, located in the intracellular D_III_-D_IV_ linker ^8-12^. Emerging evidence suggests that the IFM motif does not simply occlude the pore but rather engages in an allosteric mechanism that modulates fenestrations near the intracellular activation gate^8-10^. An alternative model proposes that hydrophobic residues in the activation gate, rather than the IFM motif alone, play a direct role in the fast inactivation process^13^, suggesting a more complex interplay between the PD and the IFM.

Previous functional studies demonstrated that the transition of the IFM motif from an unbound, solvent-exposed state to a bound state within the IFM receptor pocket is essential for activation gate closure during fast inactivation^14,15^. However, in the open state structure in which the IFM motif is mutated to QQQ, the III-IV linker including the IFM motif is unresolved^9^. In our recent full-length structure of human Na_v_1.5 in an open conformation, we identified a stabilizing interaction between the DIII-DIV linker and the EF-hand domain of the C-terminal domain (CTD), along with a loosely bound conformation of the IFM motif within the IFM receptor pocket^10^. These structural insights differ from the conventional model of IFM motif displacement which was primarily established from functional studies utilizing substituted cysteine accessibility method approaches^9,11,14,16-18^.

In this study, we investigated the gating mechanisms of human Na_v_1.5 (hNa_v_1.5) using structure-function analysis. The cryo-EM structure of full-length hNa_v_1.5 in the intermediate open state is stabilized by interactions between the N-terminal domain (NTD) and the intracellular end of the S6 segment in D_I_ (S6_I_). This structure conducts Na^+^ ions during molecular dynamics simulations. Electrophysiological measurements reveal that salt bridge interactions maintaining the IFM motif shape are critical for fast inactivation. We also identified a second Na^+^ ion binding site near the IFM motif. We propose that this binding site is also accessible to other positively charged ions such as Ag^+^, which is a common probe used as a labeling agent in cysteine accessibility assays^14^. Together, our findings reveal an alternative pathway for IFM motif accessibility by positive ions during channel opening, even when the IFM is sequestered within its canonical receptor site.

## Results

### Structure of human Na_v_1.5

To investigate the gating mechanisms of Na_v_1.5, we purified full-length protein (Extended Data Figs. 1 and 2) for cryo-EM studies. Image processing and the three-dimensional reconstruction resulted in a cryo-EM density map with a global resolution of 3.48 Å (Fig. 1 and Extended Data Fig. 2). Subsequent two rounds of local refinement further improved the EM density across most regions (Fig. 1e and Extended Data Fig. 2). The atomic model built from this map includes the transmembrane core, extracellular domains, the III-IV linker, a helical segment of the II-III linker, and the N-terminal domain (NTD) (Fig. 1a,b). Structural comparisons with the previously reported Na_v_1.5 structures revealed a dilation of the PD (Fig. 1c and Extended Data Fig. 3b,c). Notably, we identified density corresponding to a glyco-diosgenin (GDN) molecule at the activation gate (AG) region, which stabilizes the dilated conformation (Fig. 1c). This stabilization is accompanied by lateral shifts of all four S6 segments at the AG and the outward displacement of the S4-S5 linker helices (Extended Data Fig. 3c). We further observed a counterclockwise rotation of VSDs relative to the PD when viewed from the intracellular side (Extended Data Fig. 3b). Superimposition of the PDs with the partially inactivated state structure Na_v_1.5-E1784K indicated larger root mean square deviations (RMSDs) for the S6_I_, S6_II_, and S6_III_ segments at the AG region (Extended Data Fig. 3d), while the regions near the selectivity filter (SF) remained largely unchanged. These conformational differences reflect a transition from an α-helical to a π-helical configuration in the S6_I_ and S6_III_ segments. The binding of the GDN molecule stabilizes the π-helix conformation around the flexible ‘Gly-Ser’ (GS) motif of S6_I_ and S6_III_ (Fig. 1f,g), while S6_II_ and S6_IV_ maintain an α-helical conformation (Fig. 1g). The S4 segments of all VSDs are shifted outward with minimal alteration in the side-chain conformations of gating charges, resembling an activated state (Fig. 1d and Extended Data Fig. 3a).

**Fig. 1.**
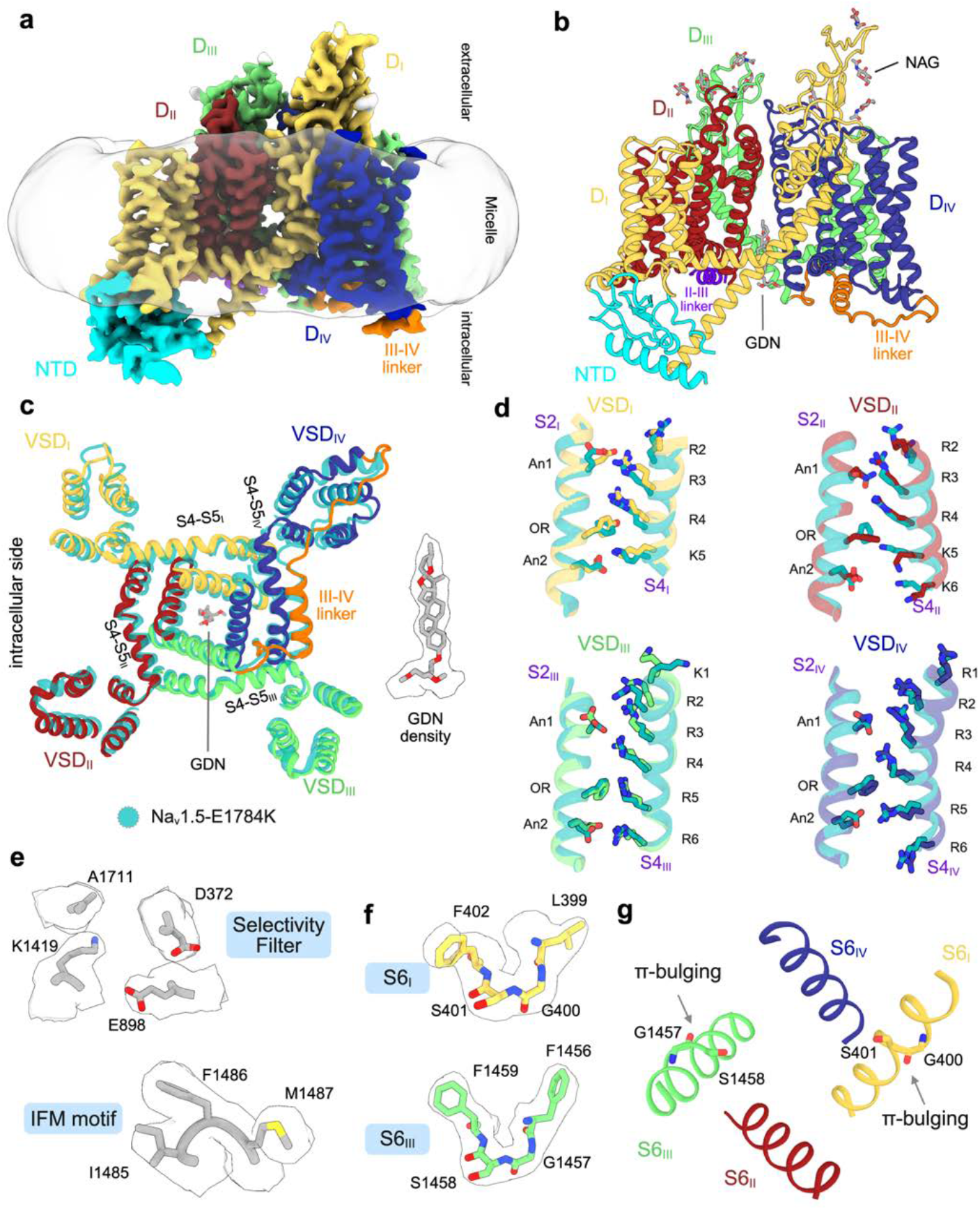
Cryo-EM structure of full-length hNa_v_1.5 in the intermediate open state. **a,** Cryo-EM map viewed from the membrane plane. Individual domains, inter-domain linkers, and detergent micelle are segmented and color-coded. **b,** Ribbon model of hNa_v_1.5 viewed from the membrane plane, with a glycol-diosgenin (GDN, grey) molecule at the AG. N-acetylglucosamine (NAG, grey) moieties are shown as sticks. All domains and interdomain linkers are segmented and color-coded according to the density map. **c,** Structural superimposition of hNa_v_1.5 with Na_v_1.5-E1784K (PDB ID: 7DTC, transparent teal) shows a lateral dilation of the overall structure. The arrangement of VSD_I_-_IV_ is shown. The density of GDN is highlighted. **d,** Comparison of S4 segments and associated gating charges (GCs) residues across VSD_I_-_IV_ between hNa_v_1.5 and Na_v_1.5-E1784K by superimposing each VSD over its S2 segment. GC residues are depicted as sticks. An1 and An2 denote anion1 and anion2, respectively. OR denotes the occluding residue. **e,** Cryo-EM densities of SF and IFM motif residues. **f,** Cryo-EM densities of the π-helical regions in segments S6_I_ and S6_III_. **g,** Ribbon representation illustrating π-bulging in segments S6_I_ and S6_III_. Segments S6_II_ and S6_IV_ remain in α-helical conformations. Residues involved in π-bulging are labeled.

### Intermediate open state conformation of the activation gate

Two hydrophobic rings located at the lower part of the S6 segments define the pore diameter at the AG, which varies depending on the kinetic state of Na_v_ channels^10^. Previous studies have reported that the diameter of an open AG is ∼10 Å to allow the passage of hydrated Na^+^ ions (7.2 Å in diameter)^9^. In our hNa_v_1.5 structure, the average pore diameters at the upper and lower layers are 10.2 Å and 10 Å, respectively (Fig. 2a and Extended Data Fig. 4a). The residues contributing to the upper layer include L409, L935, I1466, and I1768 (Fig. 2a), while those forming the lower layer are A413, L938, I1470, and I1771 (Fig. 2a). Analysis of the pore radius profile relative to other Na_v_1.5 structures revealed that the pore radius at the SF region closely matches that of other inactivated state structures (Fig. 2b and Extended Data Fig. 4b). In contrast, the region from the central cavity (CC) to the intracellular pore opening encompassing the AG, shows a pronounced dilation (Fig. 2b and Extended Data Fig. 4b). An asymmetric dilation is observed at the AG compared to structures with partially open AGs (Fig. 2b and Extended Data Fig. 3d). This dilation is more pronounced in the S6_I_, S6_II_, and S6_III_ segments than in the S6_IV_ segment (Extended Data Fig. 3d).

**Fig. 2.**
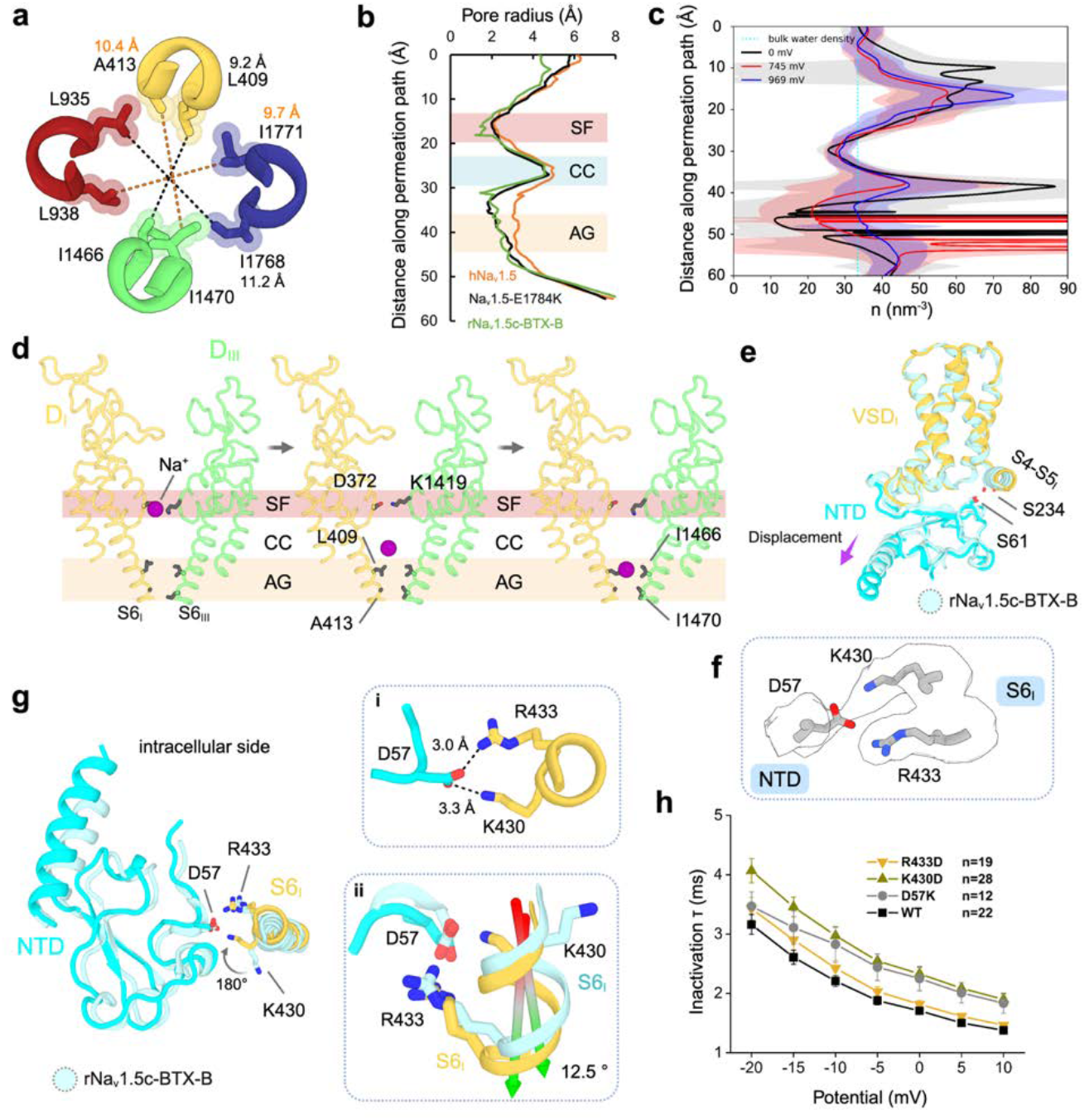
Open state conformation and NTD-S6_I_ segment interaction. **a,** Intracellular view of the activation gate formed by the S6 helices from D_1_ to D_IV_. The average diameters at the top (black dashed lines) and bottom (orange dashed lines) layers of the gate are ∼10 Å. Side chains of AG lining residues are shown as sticks. **b,** Comparison of pore radius profiles along the ion conduction pathway for hNa_v_1.5 (orange), Na_v_1.5-E1784K (black), and rNa_v_1.5c-BTX-B. (PDB ID: 7DTC, green). **c,** Time-averaged water number densities along the pore axis under transmembrane voltages (TMV) of 0 mV (black), 745 mV (red), and 969 mV (blue). At 0 mV, the activation gate region is dehydrated; hydration increases at 745 mV and becomes continuous at 969 mV. The cyan dotted line indicates bulk water density and shaded regions represent standard error. **d,** Representative frames from an MD simulation showing Na+(magenta sphere) permeating through the SF, CC, and AG. **e,** Structural alignment of hNav1.5 with rNa_v_1.5c-BTX-B (PDB: 8T6L, color: pale cyan) shows the relative position of NTD and S4-S5_I_, based on superposition of S1–S3 segments of VSD_I_. In hNa_v_1.5, the hydrogen bond between S6_1_ (NTD) and S234 (S4– S5 linker) observed in rNa_v_1.5c-BTX-B is disrupted and the NTD is displaced downward. **f,** Cryo-EM densities of residues involved in the NTD-S6_I_ interaction. **g,** Intracellular view of the interface between the NTD and the S6_I_ segment. R433 and K430 of S6_I_ are positioned near D57 of the NTD. Inset i, potential salt bridges between D57 and both R433 and K430. Inset ii, structural alignment with rNav1.5c-BTX-B reveals a 12.5° tilt and a downward shift of S6_I_ toward the NTD. **h,** Time course of inactivation (1”) from sodium current recordings show that K430D and D57K have a slower inactivation decay.

To further characterize Na^+^ conductance in our intermediate open state structure, we performed MD simulations with harmonic restraints applied to Cα positions to prevent pore collapse. In the absence of transmembrane voltage (TMV), the pore remained largely dehydrated and nonconductive near the AG (Extended Data Fig. 4c), with water density significantly lower than that of the bulk solvent. This dehydration is likely attributable to hydrophobic residues lining the AG region (Fig. 2c). Application of a TMV of 745 mV enhanced hydration at the AG likely by promoting conformational transitions toward a partially hydrated state (Extended Data Fig. 4c). The water density in this region remained lower than bulk solvent levels, suggesting the presence of substantial free energy barriers to Na^+^ permeation (Fig. 2c). At a higher TMV of 969 mV, the pore became fully hydrated and conductive (Fig. 2c, Extended Data Fig. 4c, and Supplementary Movie 1). We quantified ion conduction by monitoring cumulative permeation events over 500 ns MD trajectories. A conduction event was recorded after the complete passage of a Na^+^ ion from the extracellular side through the SF, PD, and AG to the intracellular side (Fig. 2d). During the 500 ns simulation at 969 mV, we observed three spontaneous Na^+^ conduction events (Supplementary Movie 1). In comparison, our previous open state structure demonstrated three Na^+^ permeation events at a lower TMV of 713 mV^10^. The requirement for a higher TMV in our current structure positions it between the canonical open state and the fast-inactivated state, establishing it as a bona fide intermediate open conformation in the Na_v_1.5 gating continuum.

### The S6_I_ segment - NTD interaction regulates activation gate mechanisms

The NTD of Na_v_1.5 in our structure is positioned intracellularly at the base of the VSD_I_, in agreement with other Na_v_1.5 structures (Figs 1b, 2e, and Extended Data Fig. 4d)^10,19^. A previous study identified a hydrogen bond between S62 of the NTD and S235 of the S4-S5_I_ linker as a key stabilizing interaction for the NTD conformation (Fig. 2e and Extended Data Fig. 4e,f)^19^. Structural alignment of the VSD_I_-NTD region between hNa_v_1.5 and rNa_v_1.5c-BTX-B (PDB ID: 8T6L) revealed a downward displacement of the NTD and a concomitant shift of the S4-S5_I_ linker in hNa_v_1.5 (Fig. 2e). This displacement increased the distance between the two serine residues and precluded the formation of a hydrogen bond (Fig. 2e and Extended Data Fig. 4f). In hNa_v_1.5, the stabilization of the NTD is mediated by the intracellular end of S6_I_, where D57 in the NTD forms salt bridges with K430 and R433 of S6_I_ (Fig. 2f,g). In addition, an overall dilation of the PD and a tilt of the intracellular end of S6_I_ toward the NTD were observed (Extended Data Fig. 4g,h). The helical region encompassing K430 to R433 exhibits a tilt angle of 12.5° relative to rNa_v_1.5c-BTX-B (Fig. 2g). Although the side chain conformation of R433 remains unchanged, K430 undergoes a ∼180° rotation toward D57 (Fig. 2g). To assess the functional importance of the S6_I_-NTD interaction on channel gating, site-directed mutagenesis and electrophysiological recordings were performed. Charge-reversal mutations at key interacting residues (D57K, K430D, and R433D) did not significantly alter activation, steady-state inactivation, or recovery from inactivation (Extended Data Fig. 5a,b). However, the D57K mutation resulted in reduced current density (Extended Data Fig. 5c). The D57K and K430D mutations were associated with a slowed time course of inactivation (Fig. 2h).

To dissect the conformational dynamics of the S6_I_ segment, we performed 3D variability analysis (3DVA) to resolve motions along principal components^20^. Analysis along the leading principal component revealed 14 frames that showed a swing-like motion of the intracellular end of S6_I_ (Fig. 3a and Extended Data Fig. 5d). The S6_I_ density peaked at frame 1, decreased to a minimum at frame 7, and subsequently increased through frame 14 (Supplementary Movie 2). In frame 1, the intracellular end of S6_I_ projects away from the activation gate and the pore is open (Fig. 3b,c,d and Extended Data Fig. 5e). Frame 14 shows a distinct rotation of the intracellular end of S6_I_ that results in a narrower AG (Fig. 3b,c,d and Extended Data Fig. 5d,e). Structural alignment of these models revealed a shortening of S6_I_ in frame 14 that is accompanied by a ∼13° tilt towards the center of the AG (Fig. 3d). To resolve side-chain positions, we further partitioned the principal component into two clusters and obtained reconstructions at ∼3.8 Å resolution (Extended Data Fig. 6a,b). Comparison of the maps revealed distinct intracellular positions of S6_I_ (Extended Data Fig. 6c), with an improved local resolution for S6_I_ in cluster 1 (Fig. 3e). In cluster 1, R433 forms a salt bridge with D66 of the NTD, while K430 remains distant from the NTD (Fig. 3f and Extended Data Fig. 6d). In cluster 2, both K430 and R433 engage in salt-bridge interactions with D57 of the NTD (Fig. 3f and Extended Data Fig. 6d).

**Fig. 3.**
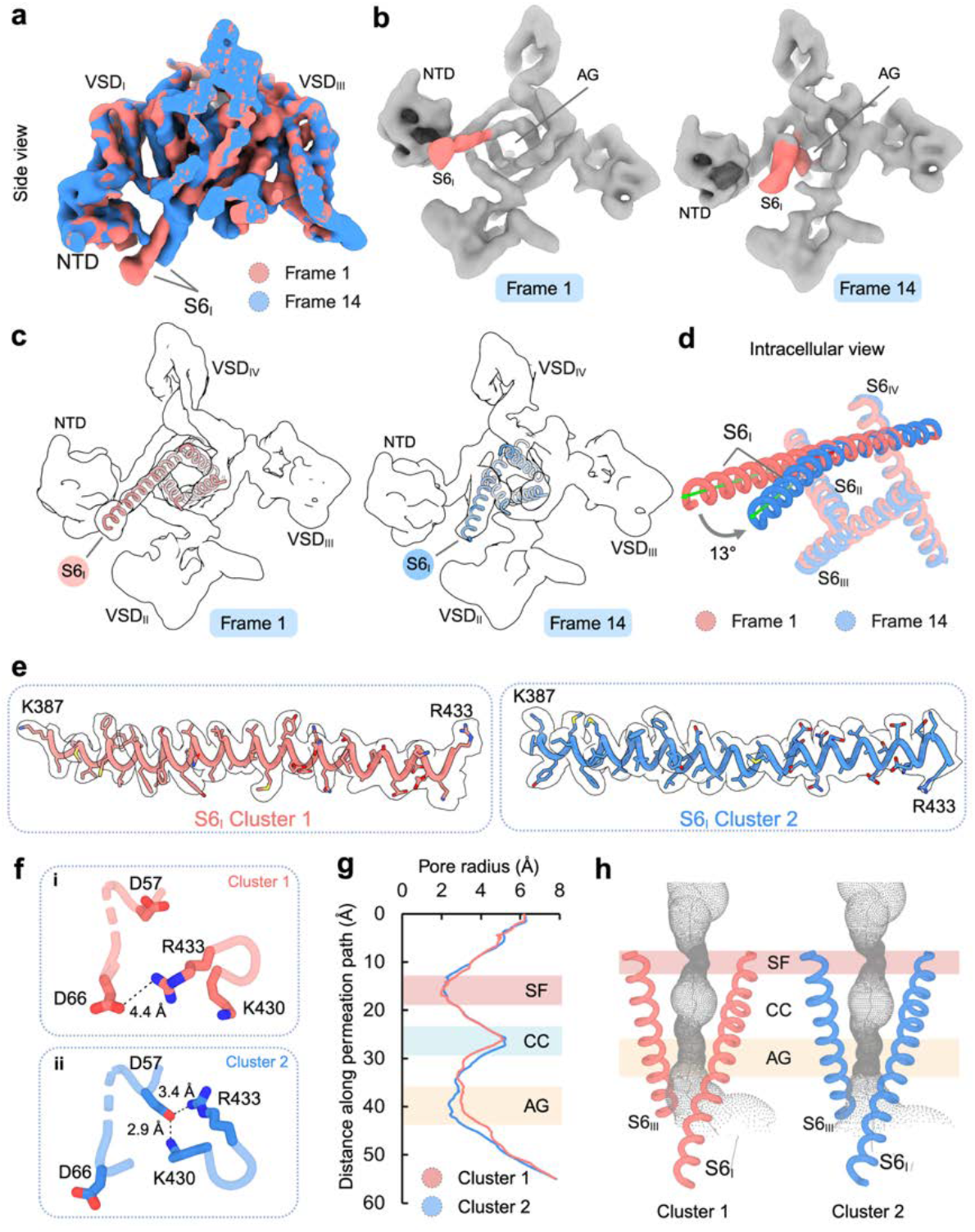
S6_I_ segment dynamics are associated with activation gate conformation. **a,** 3D variability analysis of cryo-EM map after consensus refinement and viewed from the membrane plane. The first frame map (Frame 1) and the last frame map (Frame 14) are aligned. The relative positions of S6_I_ segments density are marked. S6_I_ density in Frame 14 is shorter as compared to Frame 1. **b,** Intracellular view of Frame 1 and Frame 14. These maps highlight shift of the S6_I_ segment toward the AG which correlates with a transition from an open to a more constricted, inactivated-like conformation. **c,** The S6 segments (S6_I_-_IV_) fitted into the densities of Frame 1 and Frame 14 exhibit varying positions of S6_I_. **d,** Superimposed intracellular views reveal a 13° inward tilt of the S6_I_ segment in Frame 14 relative to Frame 1. **e,** Cryo-EM densities for the S6_I_ segments. The densities are contoured at 5σ for Cluster 1 and Cluster 2. Selected residues are labeled. **f,** NTD and S6_I_ segment interactions in Cluster 1 and Cluster 2. In Cluster 1, R433 of S6_I_ forms a salt bridge with D66 of the NTD (inset ‘**i**’). In Cluster 2, K430 and R433 of S6_I_ are positioned close to D57 of the NTD, forming a distinct salt bridge network (inset ‘ii’). **g,** Pore radius profiles along the conduction pathway for Cluster 1 (salmon) and Cluster 2 (marine) show a narrower activation gate in Cluster 2. **h,** Ion permeation paths (grey dots) overlaid with S6_I_ and S6_III_ segments. Regions corresponding to the SF, CC, and AG are indicated.

To further analyze the reduction in AG diameter observed across frames 1 to 14 in the principal component analysis, we compared the pore radius profiles between structures from cluster 1 and cluster 2 which showed significant differences between CC and AG. In cluster 2, the pore widens just below the CC but then narrows again at the AG, whereas in cluster 1, the pore remains uniformly wider (Fig. 3g,h). Residues at the AG are well resolved in both structures (Extended Data Fig. 6e). The average pore diameters at the AG are ∼10 Å in the upper layer and ∼9.8 Å in the lower layer in cluster 1 (Extended Data Fig. 6f). Cluster 2 shows reduced averaged pore diameters of ∼9.3 Å (upper layer) and ∼9.5 Å (lower layer) (Extended Data Fig. 6f). An AG diameter < 10 Å in cluster 2 suggests that it precludes the passage of hydrated Na^+^ ions.

### IFM conformation and accessibility

The positioning of the IFM motif relative to its receptor is crucial for the fast inactivation mechanisms of mammalian Na_v_ channels. The AG adopts either a closed or partially open conformation, while the IFM motif remains sequestered within its hydrophobic receptor in reported Na_v_1.5 structures^10,19,21,22^. Previous biochemical and mutagenesis studies suggest that detachment of the IFM motif from its receptor is a prerequisite for AG opening and Na^+^ permeation^14,15^. In our Na_v_1.5 structure, the IFM motif maintains its characteristic ‘U’-shaped conformation and is inserted into its receptor (Fig. 4a). Subtle repositioning of the IFM motif accompanied by a downward displacement of the adjacent short α-helix in the III-IV linker is observed (Fig. 4b). Comparative analysis of side chain conformations across available Na_v_1.5 structures reveals a conserved interaction pattern within this region. Specifically, the backbone oxygen of D1484 forms a highly conserved salt bridge with K1492, located on the short α-helix of the III-IV linker (Fig. 4c-f and Extended Data Fig. 7a). In contrast, the presence of the second salt bridge between K1492 and the backbone oxygen of M1487 within the IFM motif varies with the activation state of the channel: in Na_v_1.5-E1784K, the interaction distance is shorter (2.7 Å) compared to hNa_v_1.5 (4.4 Å) and rNa_v_1.5c-BTX-B (4.2 Å) (Fig. 4d-f and Extended Data Fig. 7a). The distance between the backbone oxygens of D1484 and M1487 remains consistent (4-5 Å), which is essential for preserving the ‘U’-shaped conformation of the IFM motif (Fig. 4f). To assess the importance of this interaction in the inactivation kinetics of hNa_v_1.5, we used site-directed mutagenesis to substitute K1492 with a negatively charged residue (K1492D). Electrophysiological measurements (Fig. 4g) showed a markedly impaired fast inactivation decay (Fig. 4h) and reduced peak current density (Fig. 4i) for K1492D, while other biophysical properties such as conductance, steady-state inactivation, and recovery from inactivation were not significantly affected (Extended Data Fig. 7b-e).

**Fig. 4.**
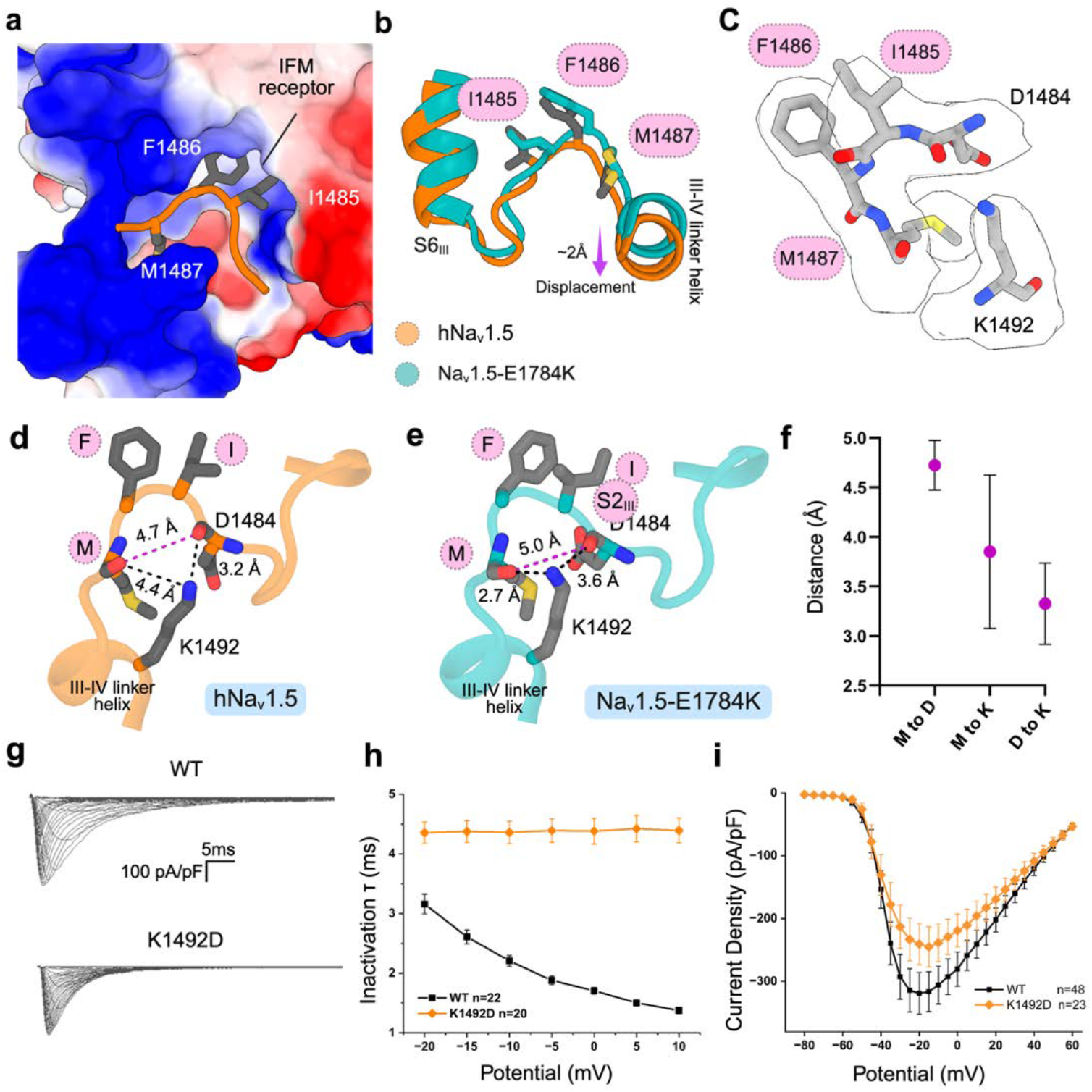
Salt bridge interactions stabilize the IFM-mediated inactivation. **a,** Electrostatic surface of the IFM receptor pocket. The side chains of I1485, F1486, and M1487 are labeled and shown as sticks. **b,** Structural superposition of hNav1.5 and Na_v_1.5-E1784K shows conformational changes in IFM residues along with a downward displacement. **c,** Cryo-EM densities of the residues involved in salt bridge interactions at the IFM motif. **d,** Backbone oxygens of D1484 and M1487 form a stabilizing salt bridge network with K1492 near the IFM motif. **e,** Reduced distance between M1487 and K1492 indicates a more stable salt bridge network at the IFM motif in the Nav1.5-E1784K structure. **f,** Variation in the distances between M1487 to D1484 (M to D), M1487 to K1492 (M to K), and D1484 to K1492 (D to K) calculated using known Na_v_1.5 structures with resolved densities. Error bars representing standard deviation (SD). **g,** Representative sodium current traces from patch-clamp recordings of K1492D. **h,** Time constants of inactivation (1”) for WT and K1492D. Results show that K1492D displayed a slowed inactivation lacking voltage dependency. **i,** Current-Voltage (I-V) relationship for WT and K1492D. The mutant resulted in decreased current density.

### Secondary Na^+^ binding site

During our structural analysis, we observed prominent density within a hydrophilic pocket adjacent to the IFM motif (Fig. 5a). This density was deduced as a Na^+^ binding site for the following reasons: First, Na^+^ ions are the only cations present throughout the purification process. Second, the density is linked to the negatively charged side chain of D1484 (Fig. 5a). Third, a similar density for Na^+^ ion has been reported for D361 at the SF region in the human Na_v_1.7-β1-β2 complex^23^. We propose that D1484 and surrounding residues act as a new binding site for Na^+^ or other monovalent cations in hNa_v_1.5. Additionally, the IFM motif is surrounded on one side by the hydrophobic IFM receptor pocket and on the other side by the hydrophilic Na^+^ ion binding pocket (Fig. 5b). We compared the positions of key residues forming the hydrophilic Na^+^binding pocket in our intermediate open state hNa_v_1.5 structure and in the partially inactivated Na_v_1.5-E1784K structure. In our structure, we observed that D1484, E1773, and Y1495 are shifted outward, forming an electronegative pocket capable of accommodating Na+(Fig. 5c). Additionally, there is a subtle repositioning of the IFM residues. The volume of this pocket measures 282 Å^3^ and exhibits two distinct regions of electrostatic surfaces (Fig. 5d). The outer region of this pocket is created by the side chains of Q1483, N1496, and K1499 (Extended Data Fig. 7f). The inner region is surrounded by D1484, F1486, M1487, Y1495, S1653, L1772, E1773, and S1776 (Extended Data Fig. 7g). As a consequence, the side chains of F1486 and M1487 in the IFM motif are positioned closer to the bound Na^+^ ion (4.7-5.1 Å), making them more accessible than I1485, which is positioned farther away at 7.2 Å (Extended Data Fig. 7h). Comparative analysis showed the presence of a significantly smaller pocket with a volume of 86 Å^3^ in Na_v_1.5-E1784K. This pocket is situated on the intracellular side of the channel, directly beneath the IFM motif (Fig. 5e). The conformation of the side chain of D1484 prevents Na^+^ coordination and binding.

**Fig. 5.**
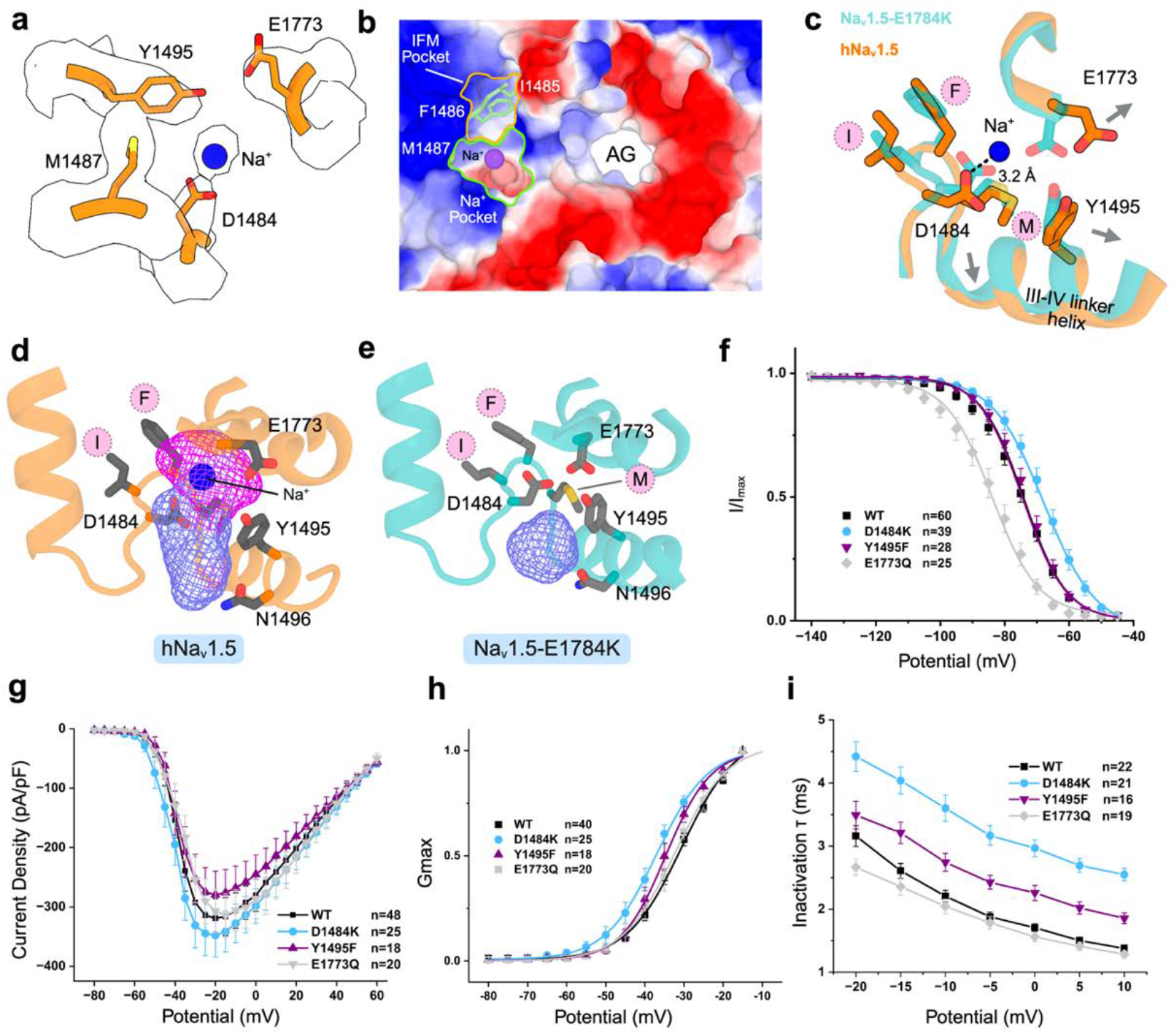
A secondary Na^+^binding site at the IFM motif region. **a,** Cryo-EM density around the IFM region of hNav1.5: a spherical density (blue) corresponds to a Na+ion coordinated by the carboxylate moiety of D1484. **b,** Electrostatic surface view of the intracellular vestibule highlighting a negatively charged Na+pocket (green outline) adjacent to the IFM pocket (yellow) and the AG. **c,** Superposition of hNav1.5 and Nav1.5-E1784K structures shows a Na^+^-D1484 interaction (3.2 Å) and a concerted displacement of E1773, Y1495 and the III-IV linker helix. **d,** Secondary binding site analysis of hNav1.5 reveals a large (282 Å^3^) electronegative Na+binding pocket near the IFM residues. The blue and magenta colors of the pocket region indicate positive and negative electrostatic potential, respectively. **e,** Secondary binding site analysis of Nav1.5-E1784K shows a markedly reduced binding pocket volume (86 Å^3^). This structure lacks the negatively charged pocket seen in WT hNav1.5. **f,** Steady-state inactivation relationship showed that E1773Q presents a left shift whereas D1484K is shifted to depolarized potentials. **g,** I-V relationship represents electrophysiological recordings from D1484K, Y1495F, and E1773Q. D1484K triggered an increase in the current density. **h,** D1484K displayed a left shift in the conductance curve in comparison to Y1495F and E1773Q. **i,** Time course inactivation relationship showed that D1484K allowed a slower inactivation whereas E1773Q has a faster inactivation.

To assess the accessibility of the Na^+^ binding pocket to silver ions (Ag^+^), which can serve as labeling agents for cysteine-substituted IFM residues^14,24^, we conducted an all-atom MD simulation under isothermal-isobaric (NPT) conditions. The system was solvated in 150 mM AgCl and subjected to a transmembrane potential of 360 mV. It is important to note that Ag^+^ is not commonly parameterized in standard biomolecular force fields. In this simulation, Ag^+^ was used to mimic NaCl conditions, rather than to specifically model Ag^+^-protein interactions. Despite the modest applied voltage, Ag^+^ permeated the Na+binding site within the first 16.5 ns of simulation (Supplementary Movie 3). The bound Ag^+^ remained coordinated in the pocket for the remainder of the 110 ns trajectory (Extended Data Fig 8a-c and Supplementary Movie 3). Structural analysis revealed that Ag^+^ is coordinated by the carboxylate side chains of D1484 and E1773 (Extended Data Fig 8d). In addition, the ion localized near the IFM motif, suggesting a possible interaction (Extended Data Fig 8a-c). Therefore, cysteine-substituted IFM motif residues are likely to form stable thiolate-Ag^+^ coordination complexes and possibly alter fast inactivation kinetics.

To understand the importance of Na^+^ coordination inside this pocket in the intermediate open state structure of hNa_v_1.5, multiple charge reversal and altered electrostatic mutations were generated for electrophysiological measurements (Extended Data Fig. 9a). Supporting the important role of D1484 in channel function, charge reversal of D1484 to a lysine (D1484K) affected several biophysical properties of the channel (Fig. 5f-i and Extended Data Fig. 9a-c). Specifically, the steady-state inactivation was shifted toward more depolarized potentials (Fig. 5f), activation was shifted toward hyperpolarized voltages (Fig. 5h), and inactivation decay was significantly slowed (Fig. 5i). Mutation of E1773 to Q1773 in the inner region of the pocket led to the stabilization of steady-state inactivation as seen by a significant hyperpolarizing shift (Fig. 5f) without affecting other biophysical properties of this mutant channel. Mutation of residue Y1495 to F1495 in this same inner region of the pocket slightly slowed the inactivation decay (Fig. 5i) without affecting any other biophysical properties of the channel. None of the mutations had a significant impact on current density (Fig. 5g) or the late sodium current (Extended Data Fig. 9c).

## Discussion

Our cryo-EM structure of wild-type hNa_v_1.5 reveals an intermediate open conformation of the activation gate, despite VSD_IV_ adopting a fully activated state and VSDs from D_I_-D_III_ appearing in an inactivated conformation. Our structure provides important structural insights into activation gate dynamics and alternative IFM accessibility.

Notably, the open state structure rNa_v_1.5c/QQQ lacks the canonical IFM motif, an essential component of fast inactivation^9^. In the inactivated structures, the IFM motif is docked into its hydrophobic receptor site, stabilizing the closed or partially closed state of the activation gate^21,22,25^. The prevailing model posits that the release of the IFM motif from this receptor is required for gate opening, a concept derived primarily from substituted cysteine accessibility assays^14^. However, structural evidence delineating conformational transitions of the IFM motif during gating remains limited and thus impedes a full understanding of the structural rearrangements required for Na^+^ conduction. Our recent wild-type hNa_v_1.5 open state structures revealed a repositioning of the IFM motif which is stabilized through interactions with the CTD^10^. In the present study, we demonstrate the regulatory influence of NTD-S6_I_ interactions on activation gate opening and propose an alternative model of IFM motif accessibility in the open state, wherein a secondary pocket near the IFM region accommodates positively charged ions.

Our cryo-EM structure of hNa_v_1.5 captures a previously uncharacterized intermediate open conformation of the AG and thus represents a critical transition state between fully open and inactivated states. This conformation is functionally relevant, as it allows Na^+^permeation during MD simulations under depolarizing voltage. Unlike fully open structures that require strong depolarization for conduction, our structure reveals a partially conductive pore stabilized without applied voltage, likely enabled by a GDN molecule trapped at the AG. This GDN molecule occupies a conserved hydrophobic site also seen in other voltage-gated sodium channel structures with wider AGs^9,23^ and likely plays a key role in mechanically holding the gate open by separating the hydrophobic residues that otherwise collapse during inactivation. The stabilization of the AG by GDN is accompanied by a substantial dilation of the PD and coordinated rearrangements across all four S6 segments. Notably, this includes α-to-π helical transitions in S6_I_ and S6_III_, consistent with emerging evidence that such transitions are conserved in voltage-gated ion channels^19,23,26,27^. Together, these findings position our structure as a functional intermediate between closed and open states, providing critical structural insights into the gating transitions that control sodium channel activation and fast inactivation.

Unlike most cryo-EM structures of Na_v_1.5 where the NTD and the intracellular end of S6_I_ are poorly resolved, our structure captures a well-ordered interaction interface between these two regions. We observed distinct salt bridges between residues in the NTD and S6_I_, suggesting that this interdomain contact plays a stabilizing role in maintaining the open AG conformation. To further explore this coupling, we performed 3D variability analysis (3DVA) which revealed changes in S6_I_ density as the AG transitions from an open to a partially closed state (Extended Data Fig. 5d,e). This resembles observations from inactivated state Na_v_ structures where the intracellular S6_I_ is often unresolved^12,28,29^. 3DVA analysis also uncovered two conformational clusters in which S6_I_ adopts alternate interactions with the NTD: R433 and K430 of S6_I_ interact with D66 in cluster 1 and with D57 in cluster 2, correlating with changes in AG diameter (Fig. 3f). These findings provide structural evidence for a dynamic coupling between S6_I_ and the NTD, implicating these salt bridges as modulators of gating transitions. While charge-reversal mutations at these sites did not significantly alter inactivation kinetics, the reduced current density observed in the D57K mutant supports a functional role for this interaction. Together, our data establish the NTD-S6_I_ interface as a critical regulatory element in activation gate stability and conformational tuning.

We observed that AG opening is coupled with subtle remodeling of the IFM motif and its binding pocket, while the motif retains a stable U-shaped conformation within its hydrophobic receptor (Fig. 4a,b). Salt bridge interactions between K1492, located on a short α-helix near the IFM motif, and the backbone oxygens of D1484 and M1487 are crucial for stabilizing this conformation. Notably, the K1492-M1487 interaction varies depending on the conformational state of the AG. A charge-reversal mutation K1492D significantly impaired fast inactivation kinetics and reduced current density (Fig. 4h,i). These findings establish that precise repositioning of the IFM motif, but not its full exposure to solvent, is critical for AG opening.

Our structure also uncovered a novel, electronegative Na+binding pocket directly adjacent to the docked IFM motif. This pocket, formed in part by D1484 and lined by F1486 and M1487, becomes accessible only when the AG is open and collapses in the inactivated state (Fig. 5d,e), rendering D1484 inaccessible to cations. This finding fundamentally challenges the prevailing interpretation of the substituted cysteine accessibility method (SCAM) that is based on the notion that IFM labeling by positively charged reagents indicates full solvent exposure following IFM displacement^14,24^. Instead, our results suggest that Ag+and MTS reagents can access substituted cysteine residues through this pocket even when the IFM remains docked, providing an alternate mechanistic explanation for prior labeling results^14,18,24^. Furthermore, the kinetic phenotype of the D1484K mutant mirrors the effects of Ag+labeling at F1486C and M1487C, reinforcing the functional importance of this pocket^14,18,24^. Together, these findings redefine our understanding of IFM accessibility, uncover a functionally coupled and structurally tuneable Na+pocket, and identify a new allosteric site with therapeutic potential for modulating inactivation in Na_v_1.5 disease causing mutations.

## Methods

### Purification of the full-length full length hNav1.5

Details about cloning expression and purification of full length hNav1.5 have been described elsewhere^10^. Briefly, the cell pellets were suspended in lysis buffer containing 25 mM HEPES (pH 7.4), 150 mM NaCl, 0.1 mM EGTA, and 10% glycerol (buffer A) including protease inhibitors. Following homogenization, the membrane was isolated through ultracentrifugation and carefully mixed in buffer A supplemented with protease inhibitors, 1% (w/v) n-dodecyl-β-D maltopyranoside (DDM, Anatrace), and 0.1% (w/v) cholesteryl hemisuccinate (CHS, Anatrace) at 4°C for 2 hours. Following ultracentrifugation, the supernatant was incubated for 2 hours with 5 mL of anti-Flag M2 affinity gel in buffer B (buffer A supplemented with 0.06% (w/v) glycol-diosgenin (GDN, Anatrace) and protease inhibitor cocktail). The column was washed with buffer B, followed by the elution of the protein using buffer B supplemented with 200 μg/ml of FLAG peptide. The eluted protein was subjected to affinity purification using Strep-Tactin XT 4Flow (IBA). Following a wash with buffer B, the bound protein was eluted using buffer B supplemented with 50 mM biotin (IBA). The eluent was concentrated and subsequently purified using a Superose 6 increase 10/300 gl column (Cytiva) in buffer C (25 mM HEPES at pH 7.4, 150 mM NaCl, 0.1mM EGTA and 0.06% GDN). The peak fractions of the purified protein were pooled and concentrated to around 8 µM for structural studies.

### Cell culture and expression of hNa_v_1.5 for electrophysiological recordings

HEK293 cells were cultivated in DMEM high glucose media (Gibco) supplemented with 10% fetal bovine serum (Gibco) and 1% penicillin-streptomycin (Sigma) at 37°C in 5% CO2. The Nav 1.5-WT and mutations plasmids were transfected into the HEK293 cells by electroporation using ATx from MaxCyte (Gaithersburg, MD) for maximal transfection efficiency according to the manufacturer’s instructions. Cells were dissociated at about 70% confluency and mixed with the target plasmid (200 ng/ µl) and transfection buffer (MaxCyte). The electroporated cells were incubated for 20 min at 37°C in 5% CO2 and then transferred to the maintenance DMEM media for about 48 hours until electrophysiological recordings.

### Whole-cell electrophysiology

Electrophysiological experiments were performed using a high-throughput automated patch-clamp with the SyncroPatch 394i (Nanion, Munich Germany). Briefly, single-hole, low-resistance recording chips from the same manufacturer were used to record sodium currents. The extracellular solution contained 140mM NaCl, 4mM KCl, 2mM CaCl_2_, 1mM MgCl2, 5mM D-glucose, 10mM HEPES. The intracellular solution contained: 10mM HEPES, 10mM EGTA, 110mM CsF, 10mM NaCl, and 10mM CsCl. Voltage protocols generation, data collection, and data analysis were performed using PatchController384 V.1.3.2 and DataController384 V1.10.1 (Nanion, Munich Germany).

The current-voltage curves of the sodium currents were elicited by holding the cells to

-120mV and stepping from -80mV to +60mV in 5mV intervals (each step held for 30ms). For activation, the G/V curve was obtained by fitting the linear part before the peak of the current-voltage curve with a Boltzmann function:

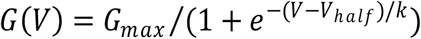

The time course of inactivation was obtained by fitting the current traces obtained from the current-voltage relationship with a single exponential fitting of each current trace from peak current to the end of the pulse (30ms) using the following equation:

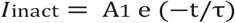

where A is the amplitude, 1” is the time constant, I is the current, and t is the time.

The recovery from inactivation was recorded with a two-pulse protocol. The pre-pulse and the test-pulse duration are 30ms, stepping from -120mV to -30mV. The interval between the two pulses ranges from 1ms to 250ms. Currents from the recovery from inactivation were fitted to the following equation:

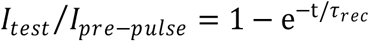

The steady-state inactivation was studied with a 500 ms pre-pulse ranging from - 140mV to -30mV, followed by a 30ms test pulse stepping from -120mV to -30mV. The currents for the steady-state inactivation were fitted to a Boltzmann distribution using the following equation:

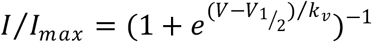

The late Na current was recorded with a 300ms pulse ranging from -120mV to -30mV. Late current was calculated from the percentage of the current measured at 250ms to the peak current.

The plots for steady-state inactivation and recovery from inactivation analysis were generated with Origin 10.1.1 software (OriginLab Corporation, Northampton, MA). The leak subtraction protocol was used.

### Statistical analysis

Statistical analyses for electrophysiology data were performed using the standard statistical package in Origin 10.1.1 (OriginLab Corporation, Northampton, MA). The student’s t-test was performed at a significance level of p < 0.05 for a single comparison after a normality test with the Shapiro-Wilk method for sample sizes 7-50. Two-sided p-values less than 0.05 were considered statistically significant. Multiple comparisons with the different mutants were performed using one-way ANOVA, and p-values less than 0.05 were considered statistically significant. Results were presented as mean ± SEM.

### Sample preparation and cryo-EM data collection

A 3.5 µL of full-length hNav1.5 protein was applied to glow-discharged UltrAuFoil 300 mesh, R1.2/1.3 grids (Quantifoil) at 4 °C and 100 % humidity. Grids were blotted for 3.5 s and then plunge-frozen in liquid ethane using Vitrobot Mark IV (Thermo Fisher). After initial screening using Glacios (Thermo Fisher Scientific), optimal grids were transferred to a Titan Krios G3i microscope (Thermo Fisher Scientific) at 300 kV equipped with a K3 direct electron detector, a BioQuantum energy filter, and a Cs image corrector. A total of 4,967 movies were collected at 81,000x magnification in super-resolution mode (pixel size: 0.4495 Å). The defocus range during collecting was kept from -0.5 to -2.5 μm. The total electron dosage per movie was 60 e^−^/Å^2^. For data processing, the summed and dose-weighted micrographs were binned to 0.899 Å/pixel. Data collection statistics are shown in Table 1.

### Image processing and 3D reconstruction

All raw movies were aligned, drift-corrected and dose-weighted in cryoSPARC v4.4.1 using Patch Motion Correction^30^, and per-micrograph defocus and CTF parameters were estimated with Patch CTF. An initial blob picking step generated a set of templates for template picking from a small subset of micrographs. The picked particles were subjected to reference-free 2D classification to yield high-quality templates. After two rounds of iterative 2D classification, a clean set of ∼1000 particles were used as a reference for Topaz training and a further two rounds of 2D classification. The extracted particles were cleaned by three rounds of iterative 2D classification. A single ab initio 3D reconstruction was generated from 281,571 particles extracted with a box size of 256 pixels. After one round of homogeneous refinement and non-uniform refinement, a 3D classification was performed to split the particles into four classes. Class 1 (75,907 particles) and Class 2 (65,347 particles) showed features for the transmembrane region as well as density for the EF-hand domain of the CTD. Using these two classes with CTD, we published the role of CTD in the Na_v_1.5 function^10^. Class-4 particles are discarded because of junk particles with empty micelles.

Class 3 map showed no CTD and we focused on this class for further analysis. All subsequent data processing steps for Class 3 were performed in cryoSPARC v4.6.2. Non-uniform refinement, along with separate global and local CTF refinements, was applied to the Class 3 map after 3D classification, yielding a global resolution of 3.7 Å from 100,809 particles. To further improve the map, three decoy classes were generated from Class 4 junk particles, specifically with empty micelles. These decoy classes were used alongside Class 3 in a round of heterogeneous refinement. Particles with clear Nav1.5 features were then re-extracted with a box size of 320 pixels and subjected to another round of non-uniform refinement, improving the global resolution to 3.57 Å with 90,155 particles.

### Model building, refinement, and validation

The model of hNa_v_1.5 was built using the sharpened map and the structure of our Class 1 map with PDB ID: 8VYJ was used as a template. The initial structure was docked into the Class 3 map using the Phenix Dock in Map tool. Subsequently, the CTD and most of the unresolved intracellular regions of the docked model were removed in Coot. Iterative model building was performed using real-space refinement in Phenix and Coot to remove the outliers and improve the model refinement statistics. The final structure refined using the Class 3 map has a total of 1258 residues. Additionally, 8 NAG, 1 GDN, and 1 sodium ion were fitted into the density. Figures were prepared with PyMOL (Schrödinger, LLC) and ChimeraX. Final figures were assembled in BioRender (www.BioRender.com). Statistics for cryo-EM model refinement are summarized in Table 1.

### 3D variability analysis (3DVA) of the EM map of hNa_v_1.5

The EM map after local refinement was subjected to 3D variability analysis (3DVA) in cryoSPARC v4.6.2^30^. A local mask was used which included all four S6 segments and the NTD. A single principal component was used for the analyses. A movie for the component was prepared with UCSF Chimera using frames generated by the 3DVA display program in a simple mode. Followed by another 3DVA display program in cluster mode to separate the frames into two clusters. The initial structure from the consensus refinement of the Class 3 model was docked into the Cluster 1 and Cluster 2 maps using the Phenix Dock in Map tool. All the ligands were removed, and iterative model building was performed using real-space refinement in Phenix and Coot.

### MD simulations

Since pore hydration is a prerequisite for conduction in sodium channels and provides _31_ more information about the functionally open state of the pore compared to the geometric radius alone, we examined the hydration profiles and conductance of NaV1.5 channel pore using classical molecular dynamics (MD) simulations in GROMACS^32^ at varying transmembrane voltage (TMV).

### System preparation for MD simulations

Using the Membrane Builder^33^ from CHARMM-GUI, the structure was embedded in a homogenous lipid bilayer of 1-palmitoyl-2-oleoyl-glycero-3-phosphocholine (POPC) molecules. Water was added to both sides of the membrane and the system was charge neutralized along with the addition of 150 mM NaCl. The periodic cell dimensions were 13.3 x 13.3 x 14.9 nm and comprised of ∼250,000 atoms. The Charmm36m all-atom force field^34^ was used to describe the interactions involving protein, lipid, and ions, while the TIP3P model^35^ was used for water. Electrostatic interactions were calculated using the particle-mesh Ewald (PME) algorithm^36^ and the bonds were constrained using the LINCS algorithm^37^.

### Equilibration MD simulations

The energy was minimized using 5000 steps of steepest descent, which was followed by canonical (NVT) equilibration at 300 K using a 2 fs time integration step with harmonic position restraints on the protein and lipid. Then, isothermal-isobaric (NPT) simulations were run using a timestep of 3 fs, Parrinello-Rahman pressure coupling^38^ at 1 bar, and temperature coupling at 300 K using velocity rescaling with a stochastic term^39^ for a total MD time duration of 50 ns such that the protein’s backbone atoms were restrained (5000 kJ mol^-1^ nm^-2^) to their initial positions. The equilibrated structure so obtained was further subjected to production MD simulations at varying TMV as described below to calculate the hydration profile of the pore.

### Production MD simulations

Varying transmembrane voltage (TMV=*E_z_L_z_*) in the form of electric field (*E_z_*) was applied along the membrane normal (*z*-direction) such that *L_z_*represents the simulation box length along the electric field direction. To prevent the pore from collapsing to a nonconducting conformation, the Cα positions of the protein were restrained by imposing harmonic potentials with force constants of 1000 kJ mol^-1^ nm^-2^. Electric fields of strength: *E_z_*= 0, 0.025, 0.05, and 0.065 V/nm were applied, which correspond to the TMV of around 0, 373, 745, and 969 mV respectively in order to translocate the sodium ions from the selectivity filter to the activation gate. Each production MD simulation was run for 500 ns using a timestep of 3 fs. Channel annotation package (CHAP)^40^ was employed for the hydration profile calculations such that the initial 100 ns data was discarded, and the averages were computed over the frames extracted every 1.5 ns.

## Data availability

The cryo-EM structure is deposited in the Protein Data Bank (PDB) with a PDB ID: XXXX and Electron Microscopy Data Bank (EMDB) under the EMDB ID: XXXX. Data supporting the findings of this study are available in the article and its Supplementary information.

## Acknowledgments

Electron microscopy data were acquired at the Center for Electron Microscopy and Analysis at The Ohio State University. We thank the Ohio Supercomputer Center for high-performance computing resources. This work was supported by National Institutes of Health grant R01HL094450 (I.D.) and the Frick Center for Heart Failure via a Synergy Award from the Dorothy M. Davis Heart and Lung Research Institute at

The Ohio State University Wexner Medical Center (I.D) and start-up funds from The Ohio State University College of Medicine (S.M.H, I.D, and K.C).

## Author contributions

I.D. and K.C. designed research; R.B., A.L.L.-S., A.P., A.R.-N., and H.-L.H. performed research; A.R.-N., H.-L.H., X.C., S.M.H., I.D., and K.C. contributed new reagents/analytic tools; R.B., A.L.L.-S., A.P., and K.C. analyzed data; and R.B., A.L.L.-S., A.P., X.C., S.M.H., I.D., and K.C. wrote the paper.

## Notes

### Competing Interest Statement

The authors have declared no competing interest.

## References

1. Fozzard, H.A. & Hanck, D.A. Structure and function of voltage-dependent sodium channels: comparison of brain II and cardiac isoforms. Physiol Rev 76, 887–926 (1996).

2. Gamal El-Din, T.M. When the Gates Swing Open Only: Arrhythmia Mutations That Target the Fast Inactivation Gate of Na(v)1.5. Cells 11(2022).

3. Goldin, A.L. Mechanisms of sodium channel inactivation. Curr Opin Neurobiol 13, 284–90 (2003).

4. Ulbricht, W. Sodium channel inactivation: molecular determinants and modulation. Physiol Rev 85, 1271–301 (2005).

5. Remme, C.A. & Bezzina, C.R. Sodium channel (dys)function and cardiac arrhythmias. Cardiovasc Ther 28, 287–94 (2010).

6. Balser, J.R. The cardiac sodium channel: gating function and molecular pharmacology. J Mol Cell Cardiol 33, 599–613 (2001).

7. Shah, M., Akar, F.G. & Tomaselli, G.F. Molecular basis of arrhythmias. Circulation 112, 2517–29 (2005).

8. Wisedchaisri, G. et al. Resting-State Structure and Gating Mechanism of a Voltage-Gated Sodium Channel. Cell 178, 993–1003 e12 (2019).

9. Jiang, D. et al. Open-state structure and pore gating mechanism of the cardiac sodium channel. Cell 184, 5151–5162 e11 (2021).

10. Biswas, R. et al. Structural basis of human Na(v)1.5 gating mechanisms. Proc Natl Acad Sci U S A 122, e2416181122 (2025).

11. West, J.W. et al. A cluster of hydrophobic amino acid residues required for fast Na(+)-channel inactivation. Proc Natl Acad Sci U S A 89, 10910–4 (1992).

12. Pan, X. et al. Structure of the human voltage-gated sodium channel Na(v)1.4 in complex with beta1. Science 362(2018).

13. Liu, Y., Bassetto, C.A.Z., Jr., Pinto, B.I. & Bezanilla, F. A mechanistic reinterpretation of fast inactivation in voltage-gated Na(+) channels. Nat Commun 14, 5072 (2023).

14. Kellenberger, S., Scheuer, T. & Catterall, W.A. Movement of the Na+ channel inactivation gate during inactivation. J Biol Chem 271, 30971–9 (1996).

15. Kellenberger, S., West, J.W., Catterall, W.A. & Scheuer, T. Molecular analysis of potential hinge residues in the inactivation gate of brain type IIA Na+ channels. J Gen Physiol 109, 607–17 (1997).

16. Karlin, A. & Akabas, M.H. Substituted-cysteine accessibility method. Methods Enzymol 293, 123–45 (1998).

17. Tomaselli, G.F. Cysteine mutagenesis in the voltage-dependent sodium channel structural insights and implications. Trends Cardiovasc Med 7, 211–8 (1997).

18. Kellenberger, S., West, J.W., Scheuer, T. & Catterall, W.A. Molecular analysis of the putative inactivation particle in the inactivation gate of brain type IIA Na+ channels. J Gen Physiol 109, 589–605 (1997).

19. Tonggu, L. et al. Dual receptor-sites reveal the structural basis for hyperactivation of sodium channels by poison-dart toxin batrachotoxin. Nat Commun 15, 2306 (2024).

20. Punjani, A. & Fleet, D.J. 3D variability analysis: Resolving continuous flexibility and discrete heterogeneity from single particle cryo-EM. J Struct Biol 213, 107702 (2021).

21. Li, Z. et al. Structure of human Na(v)1.5 reveals the fast inactivation-related segments as a mutational hotspot for the long QT syndrome. Proc Natl Acad Sci U S A 118(2021).

22. Jiang, D. et al. Structure of the Cardiac Sodium Channel. Cell 180, 122–134 e10 (2020).

23. Huang, G. et al. High-resolution structures of human Na(v)1.7 reveal gating modulation through alpha-pi helical transition of S6(IV). Cell Rep 39, 110735 (2022).

24. Deschenes, I., Trottier, E. & Chahine, M. Cysteine scanning analysis of the IFM cluster in the inactivation gate of a human heart sodium channel. Cardiovasc Res 42, 521–9 (1999).

25. Li, Z. et al. Structural Basis for Pore Blockade of the Human Cardiac Sodium Channel Na(v) 1.5 by the Antiarrhythmic Drug Quinidine*. Angew Chem Int Ed Engl 60, 11474–11480 (2021).

26. Su, Q. et al. Cryo-EM structure of the polycystic kidney disease-like channel PKD2L1. Nat Commun 9, 1192 (2018).

27. Zhao, Y. et al. Molecular Basis for Ligand Modulation of a Mammalian Voltage-Gated Ca(2+) Channel. Cell 177, 1495–1506 e12 (2019).

28. Shen, H., Liu, D., Wu, K., Lei, J. & Yan, N. Structures of human Na(v)1.7 channel in complex with auxiliary subunits and animal toxins. Science 363, 1303–1308 (2019).

29. Wu, Q. et al. Structural mapping of Na(v)1.7 antagonists. Nat Commun 14, 3224 (2023).

30. Punjani, A., Rubinstein, J.L., Fleet, D.J. & Brubaker, M.A. cryoSPARC: algorithms for rapid unsupervised cryo-EM structure determination. Nat Methods 14, 290–296 (2017).

31. Jo, S., Kim, T., Iyer, V.G. & Im, W. CHARMM-GUI: a web-based graphical user interface for CHARMM. J Comput Chem 29, 1859–65 (2008).

32. Bauer, P., Hess, B. & Lindahl, E. GROMACS 2022 Manual. doi.org/10.5281/zenodo.6103568 (2022).

33. Lee, J. et al. CHARMM-GUI Membrane Builder for Complex Biological Membrane Simulations with Glycolipids and Lipoglycans. J Chem Theory Comput 15, 775–786 (2019).

34. Huang, J. et al. CHARMM36m: an improved force field for folded and intrinsically disordered proteins. Nat Methods 14, 71–73 (2017).

35. Jorgensen, W.L., Chandrasekhar, J., Madura, J.D., Impey, R.W. & Klein, M.L. Comparison of simple potential functions for simulating liquid water. The Journal of Chemical Physics 79, 926–935 (1983).

36. Essmann, U. et al. A smooth particle mesh Ewald method. The Journal of Chemical Physics 103, 8577–8593 (1995).

37. Hess, B., Bekker, H., Berendsen, H.J.C. & Fraaije, J.G.E.M. LINCS: A linear constraint solver for molecular simulations. Journal of Computational Chemistry 18, 1463–1472 (1997).

38. Parrinello, M. & Rahman, A. Polymorphic transitions in single crystals: A new molecular dynamics method. Journal of Applied Physics 52, 7182–7190 (1981).

39. Bussi, G., Donadio, D. & Parrinello, M. Canonical sampling through velocity rescaling. The Journal of Chemical Physics 126(2007).

40. Klesse, G., Rao, S., Sansom, M.S.P. & Tucker, S.J. CHAP: A Versatile Tool for the Structural and Functional Annotation of Ion Channel Pores. J Mol Biol 431, 3353–3365 (2019).

